# Dynamic up- and down-regulation of the default (DMN) and extrinsic (EMN) mode networks during alternating task-on and task-off periods

**DOI:** 10.1101/658757

**Authors:** Kenneth Hugdahl, Justyna Beresniewicz, Katarzyna Kazimierczak, Kristiina Kompus, Rene Westerhausen, Lars Ersland, Renate Grüner, Karsten Specht

**Affiliations:** Department of Biological and Medical Psychology, University of Bergen, Norway; Division of Psychiatry, Haukeland University Hospital, Bergen, Norway; Department of Radiology, Haukeland University Hospital, Bergen, Norway; Institute of Psychology, University of Oslo, Norway; Department of Clinical Engineering, Haukeland University Hospital, Bergen, Norway; Department of Education, UiT/The Arctic University of Norway, Tromsø, Norway; Mohn Medical Imaging and Visualization Centre, Haukeland University Hospital, Bergen, Norway

**Author notes:** **Address all correspondence** to Kenneth Hugdahl, Department of Biological and Medical Psychology, University of Bergen, Norway.

**Keywords:** Cortical networks, connectivity, default mode network, DMN, extrinsic mode network, EMN, auditory network, salience network

## Abstract

Using fMRI, Hugdahl et al. (2015) reported the existence of a general-domain cortical network during active task-processing which was non-specific to the cognitive task being processed. They labelled this network the extrinsic mode network (EMN). The EMN would be predicted to be negatively, or anti-correlated with the classic default mode network (DMN), typically observed during periods of rest, such that while the EMN should be down-regulated and the DMN up-regulated in the absence of demands for task-processing, the reverse should occur when demands change from resting to task-processing. This would require alternating periods of task-processing and resting, and analyzing data continuously when demands change from active to passive periods and vice versa. We were particularly interested in how the networks interact in the critical transition points between conditions. For this purpose we used an auditory task with multiple cognitive demands in a standard fMRI block-design. Task-present (ON) blocks were alternated with an equal number of task-absent, or rest (OFF) blocks to capture network dynamics across time and changing environmental demands. To achieve this, we specified the onset of each block, and used a finite-impulse response function (FIR) as basis function for estimation of the fMRI-BOLD response. During active (ON) blocks, the results showed an initial rapid onset of activity in the EMN network, which remained throughout the period, and faded away during the first scan of the OFF-block. During OFF blocks, activity in the DMN network showed an initial time-lag where neither the EMN nor the DMN was active, after which the DMN was up-regulated. Studying network dynamics in alternating passive and active periods may provide new insights into brain network interaction and regulation.

## Introduction

Applying an inclusive conjunction analysis to fMRI data from nine different studies with a total of 187 subjects, and comprising nine different cognitive tasks, Hugdahl et al. [1] found a generalized cortical network that was independent of the specifics of the individual task and cognitive processes. The authors labelled this the Extrinsic Mode Network (EMN), as a task non-specific network, with a fronto-temporo-parietal distribution, including the inferior and middle frontal gyri, inferior parietal lobule, supplementary motor area, and the inferior temporal gyrus. The EMN thus has a spatial architecture overlapping with what Fedorenko et al. [2] labelled the “cognitive flexibility network”, and Duncan [3] labelled the “multiple demand network” (see also [4]). Common for all these networks is that they are suggested to be general-domain networks, i.e. they show up-regulation of activity across a range of cognitive tasks, e.g. [5], [6], [7], [8]. The general-domain networks further share characteristics with several domain-specific networks, such as the dorsal attention and salience networks [9], [10], [11], [12]), central executive network [13], [14], fronto-parietal network [15], and ventral attention network [16], see also Lee et al. [17], and Cabeza and Nyberg [18] for overviews. In the current study, we asked how the EMN as a general-domain network relates to the more known default mode network (DMN) [19], [20], [12], see also [21], [22], [23], [24], which is typically observed in the absence of specific tasks. It could be predicted that the DMN should be negatively, or anti-correlated with a general-domain network, like the EMN. This prediction is derived from previous findings that neuronal activity varies reciprocally between characteristic DMN- and task-positive network-areas, when studied under both resting-state [16], [25], [26], [23], and during active task conditions [27], [28], [29]. Lustig et al. [27] used alternating blocks of passive viewing of a fixation cross and active processing of a semantic judgement task. Results showed increased activations in the left frontal cortex during task-processing and deactivation in the same area during passive fixation blocks. An opposite pattern was seen in the lateral parietal cortex, with deactivations during task processing. The study by Lustig et al. [27] therefore showed inverse activations and deactivations in brain areas linked to active task-processing compared to passive viewing. Following previous findings we therefore asked whether a similar relationship should hold for general-domain networks, and in particular for the EMN, when different cognitive tasks are alternated during the scanning session. Thus, we alternated brief periods with task presentations with brief periods of resting with no tasks present.

Conventional analysis of block-design data is to subtract activity during OFF-blocks from activity during ON-blocks, where the OFF-blocks act as a baseline control-condition (we leave out here the discussion in the literature whether the assumption of “pure insertions” in block-designs is a valid assumption or not, cf. [30]. The resulting activity pattern would thus reflect active task-processing. By subtracting activity obtained during task-processing ON-blocks from activity obtained during resting OFF-blocks, cf. [19], it should be possible to display activity that would be deactivated during task-processing blocks. We are not focusing here on whether task-positive and task-negative networks are anti-correlated per se in resting-state fMRI situations, cf. [31], [27], but on the time-dynamics of up- and down-regulations across the transitions between task-present and task-absent periods, with a focus on the interaction between the DMN and EMN networks. For this purpose, we used a finite-impulse response (FIR) function to model the BOLD response as implemented in the Statistical Parametric Mapping (SPM) analysis software (https://www.fil.ion.ucl.ac.uk/spm/). We re-analyzed fMRI data from >100 healthy individuals from a previous study in our laboratory [32], where the subjects had been tested with an auditory dichotic listening (DL) task with three instruction conditions that emphasized either perception [33], attention/vigilance [34] or executive control functions [35]. We chose this task because it reflects the changing cognitive demands and coping situations during an ordinary day, including both low-, perception, and high-, executive control, level demands. The so called forced-attention DL paradigm was originally developed by Hugdahl and Andersson [34] for the study of the role of cognitive factors in auditory perception, and it has repeatedly been shown to produce valid and reliable results with regard to perception, attention and executive control, see e.g. [36], [37], [38], [39], [40], [41], [42], [43]. We now report how the EMN interact with the DMN within a single paradigm which included alternating task-presence (ON) and task-absence (OFF) periods, and with recurring and varying cognitive tasks and demands. Such an experimental design would be a novel way of capturing the dynamic interaction between passive rest and active processing periods, going beyond a fMRI “resting period” data acquisition approach. In order to capture the dynamics of network up- and down-regulation over time, and especially at the transition points between ON- and OFF-blocks, we specified the onset of each condition and used a 3s finite-impulse response function (FIR) as basis function for estimation of the BOLD response. This approach modelled each scan per task-ON- and task-OFF-block separately, which would allow an analysis of the fine-grained dynamics in the critical time-window when the situation switched from active to passive, and from passive to active time periods.

## Materials and Methods

### Participants

The participants were 104 healthy adults, with mean age of 29.3 years, standard deviation 8.3 years. Approximately half of the participants were males, and half were females. The participants volunteered for participation, and further details can be found in [33]. The study was conducted according to the Declaration of Helsinki regarding ethical standards. The re-analyzed data had in addition previously been approved by the Regional Ethics Committee for Medical Research in the Western Health Region of Norway (REK-Vest), and also been completely anonymized before the current analyses were made.

### Cognitive tasks

The cognitive task was an auditory speech perception task, with repeated dichotic presentations of two different consonant-vowel (CV) syllables presented on each trial, one in the right ear and the other at the same time in the left ear. The participant is not told that there are two different sounds, one in each ear, on every trial. The baseline instruction to the participant was to report which syllable they perceived most clearly on each trial, emphasizing a single response, and with no instruction about allocation of attention to either the right or left ear. The task consists of repeated presentations of syllable-pair, using all combinations of the six stop-consonants /b/, /d/, /g/, /p/, /t/, /k/ paired with the vowel /a/, making up the CV-syllables /ba/, /ga/, /pa/, /da/, /ka/, /ta/. A trial could thus be the presentation of /ba/ in the left ear and simultaneously the syllable /pa/ in the right ear, see [36] for an overview of the dichotic listening task. The dichotic CV-syllable task has historically been used for the study of hemispheric asymmetry, which is reflected in the typical higher accuracy scores for reporting of the right ear stimulus, called a right-ear advantage (REA) [44], [35], [45], [46]. The paradigms has however also been used for the study of higher cognitive functions, like attention and executive control functions [47], [43], [48], [49], which Hugdahl and Andersson [33] labelled “the forced-attention” dichotic listening paradigm. In the latter case instructions to explicitly focus attention to and report from only the right or left ear is alternated between trial-blocks, and mixed with blocks of no-attention-focus instruction. A methodological advantage with the forced-attention variant is that it allows for the study of perception, attention and executive functions within the same experimental paradigm, by simply changing the instructions to the participants in the course of the experimental session, see [35] for further examples. In the present fMRI-variant of the DL paradigm, each of the three instruction blocks (no instruction, instruction to focus on the right ear, instruction to focus on the left ear) were repeated three times during the session. The order of the presentation of the three conditions were pseudorandomized among the non-forced (NF), forced-right ear (FR), and forced-left ear (FL) instruction epochs. To approximate an every-day situation situation with brief processing and resting periods, the nine task-present epochs (ON-blocks) were alternated with nine resting epochs (OFF-blocks) with no stimuli or instruction present. Each ON- and OFF-block had a duration of 55 sec. Since the focus of the present study was the dynamic interaction of the EMN and DMN networks, we present data averaged across the three instruction conditions, since this will capture the conglomerate activity across the three cognitive tasks, to act as a proxy of for the fluctuations of cognitive demands experienced during the course of a day.

### MR imaging

The MR scanner was a 3T GE SignaHDx scanner, and for the initial anatomical scanning, a T1 3D Fast Spoiled Gradient Recall sequence (FSPGR) was applied: TE = 14 ms, TR = 400 ms, TI = 500 ms), with 188 consecutive sagittal slices (1 mm thick, no gap, scan matrix: 256 × 256; FOV 256 mm). For the following echo-planar functional imaging (EPI), a sparse-sampling sequence was applied with TR of 5.5 sec, and with acquisition time (TA) of 1.5 sec, leaving a silent gap of 4 sec when the stimuli were presented, see [46]. 180 EPI-volumes were acquired, consisting of 25 axial slices in each volume (FOV: 220 mm; scan matrix 64 × 64; 5 mm slice thickness, 0.5 mm gap; TE = 30 ms). There were 10 EPI-volume acquisitions, or scans, during each of the task-present and task-absent blocks, each of 55 sec. Total session time for the EPI-imaging part was thus (9 × 55) × 2 = 16.5 min, with regularly alternating task-present and task-absent blocks.

### Statistical analysis and visualization of fMRI data

The fMRI-data were analyzed with the SPM12 software package ((Wellcome Department of Cognitive Neurology, London, UK, http://www.fil.ion.ucl.ac.uk/spm/), following standard SPM settings. In brief, the raw DICOM images were converted to nifty-format, and pre-processed following SPM implanted routines for realignment and unwarping, normalizing the EPI-images to the MNI template, and smoothing with an 8 mm kernel. Thereafter, 1st-level analysis was performed, by specifying the onset of each condition and using a finite-impulse response function (FIR) as basis function, which modeled each scan per ON and OFF block separately, but averaged across repetitions of the same condition. The resulting beta-images/time-bins (TBs) (20 per condition, NF, FR, FL) were then used as input for the next, 2nd-level analysis, which was defined as a 3 × 20 repeated measure ANOVA model, with the factor condition (NF, FR, FL) and the factor time-bin (TB) (1-20). This model allows exploring not only averaged bock-effects, mimicking “classical” ON-OFF contrasts, but also the temporal dynamics and time derivatives by specifying contrasts for each TB separately. (see Figure 1).

**Figure 1:**
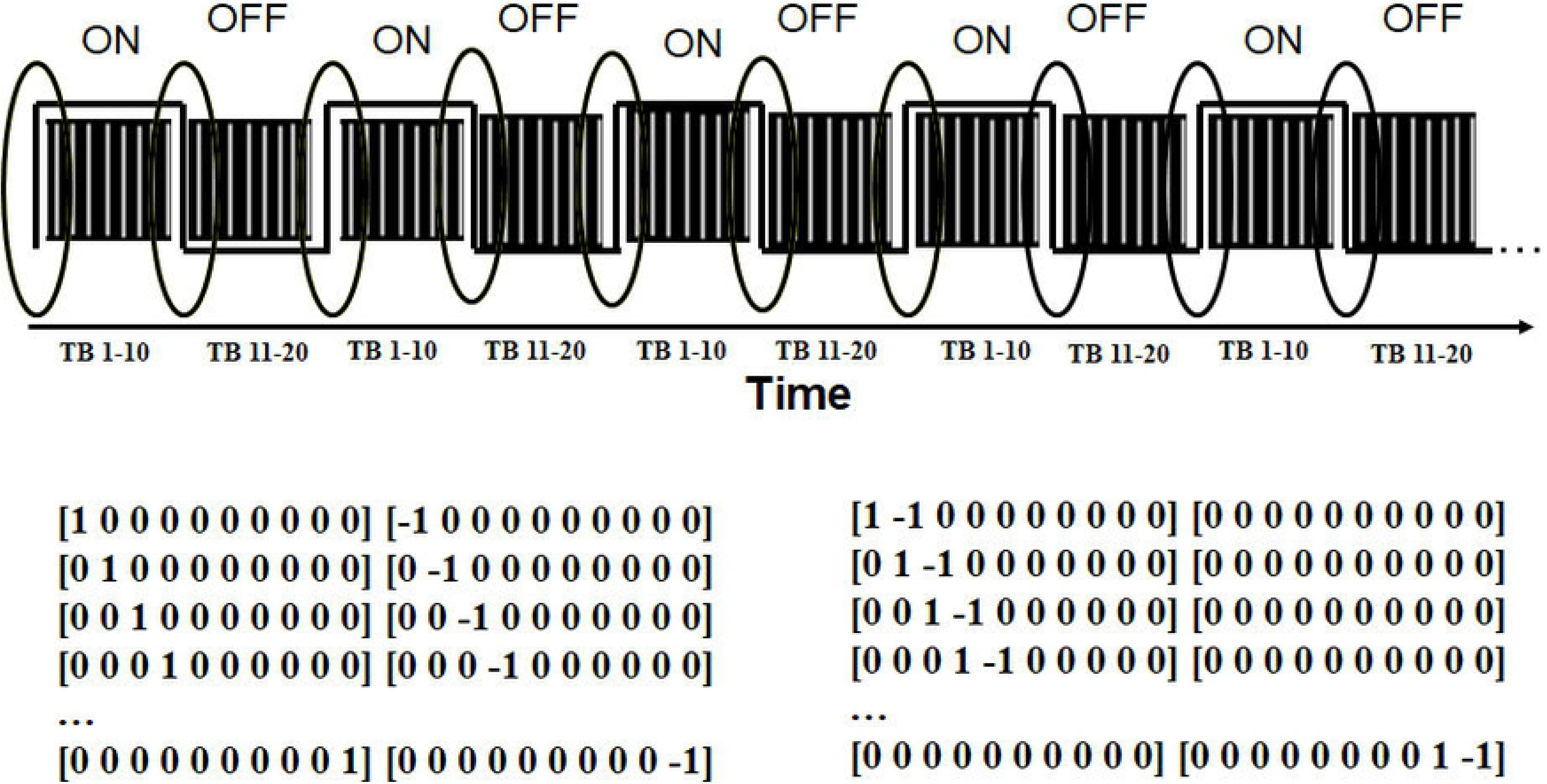
Outline of the block-design with alternating ON-and OFF-periods with corresponding task-processing and resting, respectively. The time-line illustrates the specification of the separate time-bins (TB) for respective ON- and OFF-blocks, with the transition points between blocks marked with circles. The tables under the design illustrates the contrasts used in the analysis for the time-bin (left) and time-derivative (right) analyses, respectively (see text for further details.

The following contrasts were specified: First, the average ON-OFF and OFF-ON contrasts were specified. Second, corresponding averaged TBs for the ON and OFF contrast were compared, i.e. TB1 ON against TB1 OFF, TB2 ON against TB2 OFF, etc, and repeated for the remaining eight pairs of time-bins, and t-contrasts were specified in both directions. Third, we explored the time-derivatives of the activity by contrasting averaged time-bins next to each other, i.e. TB1 against TB2, TB2 against TB3, etc, i.e., TB *n* against TB *n*+*1*. These latter contrasts will reveal consecutive significant changes from one TB to the next, i.e. from TB *n* to TB *n*+*1* as a “sliding window”. The “sliding window” contrast would be sensitive for picking up the time dynamics in the relative up- and down-regulation of the task-positive and task-negative networks, i.e. sensitive to the transition from the last TB for an ON-block to the first TB for an OFF-block, and vice versa (see Figure 1). If an activity remained unchanged from one TB to the next (as one would expect for adjacent TBs in the middle of a block), this contrast would not show anything. Again, both directions of contrasts were estimated, i.e. whether there was a significant increase or decrease from one TB to the next. For simplicity, differential effects between the three conditions (NF, FR, FL) were not explored, since the focus was on activity that were common across diverse cognitive tasks, cfr. [4], [2], [1]. Results were explored for statistical significance, using a family-wise error (FWE)-corrected threshold level of p < .05 in the main analyses, to protect against Type-I errors, and with at least 10 voxel per cluster.

## Results

The results of the averaged ON-OFF contrast revealed the typical task-positive activity pattern with bilateral activity in the auditory cortex and surroundings, and of the task-positive, EMN, network [1] in the prefrontal cortex, including the anterior and middle cingulate cortex, supplementary motor area (SMA/preSMA), and thalamus. The opposite contrast revealed areas, belonging to the task-negative, DMN, network [20], revealed significant activity in the precuneus, inferior parietal, and medial orbitofrontal areas, and in occipital areas (see Figure 2b and Table 1b). This could be activity returning to baseline during resting periods, as previously found, see e.g. [20], or as a new finding with increases above baseline in situations with alternating task-negative and task-positive periods.

**Figure 2 (a,b):**
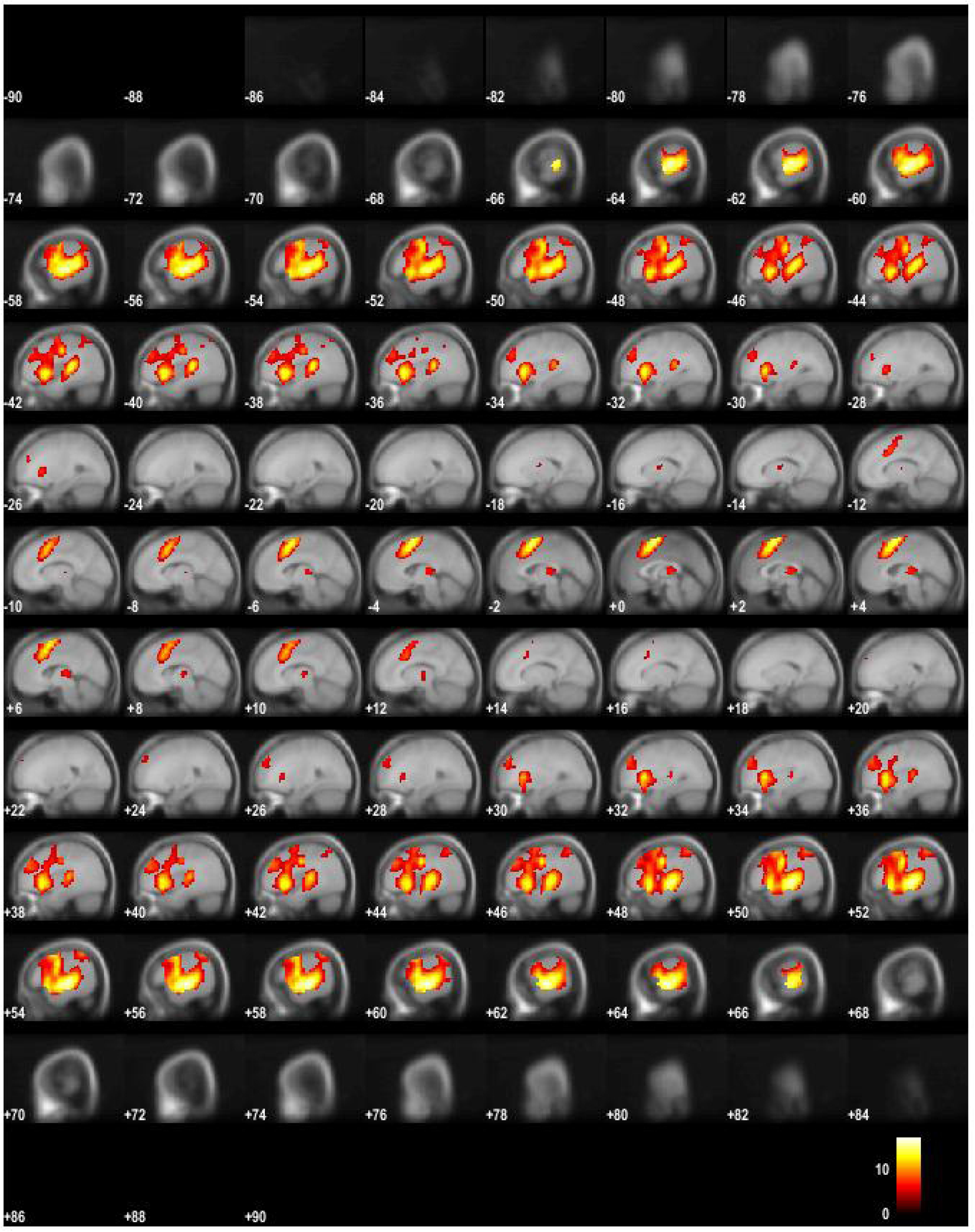
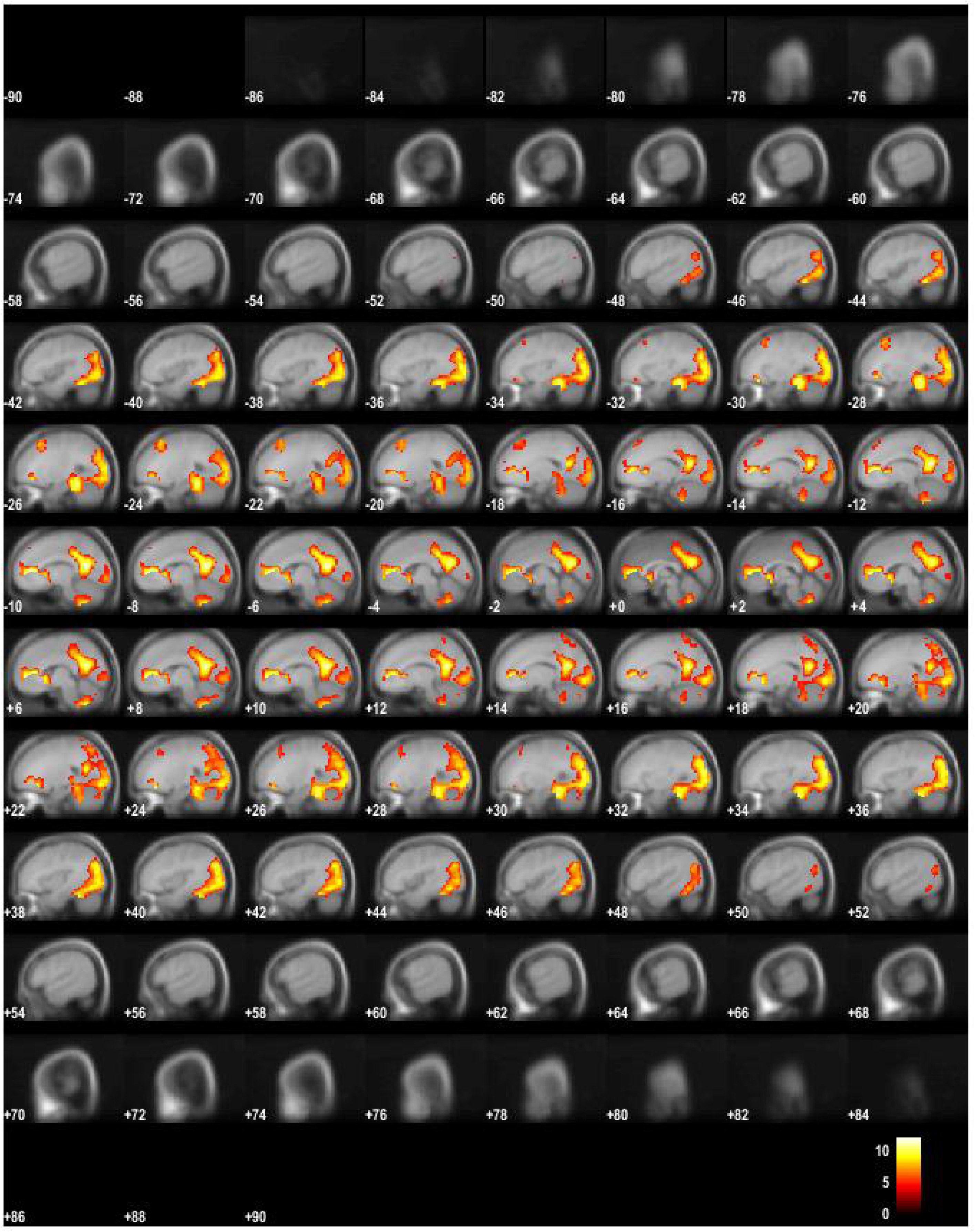
Figure 2a shows BOLD activity for the ON – OFF contrast, thresholded at FEW .05, 10 voxels, in sagittal images through the entire brain volume, from “ear-to-ear”, rendered on an average MNI template. Figure 2b shows BOLD activity for the OFF-ON OFF contrast, thresholded at FEW .05, 10 voxels, in sagittal images through the entire brain volume, from “ear-to-ear”, rendered on an average MNI template.

**Table 1a.** Summary of significantly activated clusters (with local maxima), and peak voxel x, y, z coordinates and corresponding t- and z-values and AAL atlas anatomical localizations, for the OFF-ON contrast

| Cluster size | Peak t-value | Peak z-value | X | Y | Z | Anatomical localization |
| --- | --- | --- | --- | --- | --- | --- |
| 4356 | 17.07 | Inf | 60 | -21 | -3 | rSTG |
|  | 15.78 | Inf | 57 | -12 | -6 | rSTG |
|  | 14.02 | Inf | 45 | 15 | -6 | rInsula |
| 4145 | 16.48 | Inf | -60 | -15 | 0 | lSTG |
|  | 16.44 | Inf | -60 | -24 | 3 | lSTG |
|  | 14.76 | Inf | -60 | -39 | 9 | lSTG |
| 917 | 15.11 | Inf | 0 | 6 | 60 | l/rSMA |
|  | 13.31 | Inf | 3 | 18 | 47 | rSMA |
| 119 | 6.85 | 6.78 | 0 | -27 | 6 | lThlmus |
|  | 4.90 | 4.87 | -15 | -6 | 12 | lThlmus |

**Table 1b.** Summary of significantly activated clusters (with local maxima), and peak voxel x, y, z coordinates and corresponding t- and z-values and AAL atlas anatomical localizations, for the ON-OFF contrast

| Cluster size | Peak t-value | Peak z-value | X | Y | Z | Anatomical localization |
| --- | --- | --- | --- | --- | --- | --- |
| 7669 | 12.18 | inf | -9 | -57 | 15 | lPrecun |
|  | 11.73 | inf | 12 | -54 | 18 | rPrecun |
|  | 11.31 | inf | -3 | -63 | 21 | lPrecun |
| 972 | 11.55 | inf | 6 | 51 | 0 | rSFG |
|  | 11.51 | inf | -9 | 45 | 6 | lACC |
|  | 10.93 | Inf | 12 | 39 | 6 | lSFG |
| 203 | 8.44 | Inf | -24 | 24 | 45 | lSFG |
| 40 | 5.56 | 5.52 | 27 | 27 | 45 | rMFG |

The differential, TB-wise contrasts are presented as activity changes rather than contrast-by-contrast activity. First, the temporal evolution of the task-positive networks, including the EMN network, was explored. As can be seen from Figure 3, the task-positive activity started with a strong visual activity, reflecting the on-screen instruction, followed by activity of the auditory and EMN network after about 5.5 sec (one time bin), which remained constant throughout the entire ON period and faded away during the first scan of the OFF block. Interestingly, the task-negative and DMN networks showed more dynamic changes than the task-positive and EMN networks. There was an initial time-lag of about 5.5 sec where neither the EMN nor the DMN or any other task-related networks was active. After this initial period, the DMN showed the strongest recurrence, which however faded away towards the end of the OFF block. The only activity that later evolved during the OFF blocks and remained throughout the block was the orbitofrontal recurrence. Figure 3 shows the overall activity including also line plots for the activity profiles from the posterior cingulate cortex and the right inferior frontal gyrus, representing hubs of the DMN and EMN networks, respectively.

**Figure 3:**
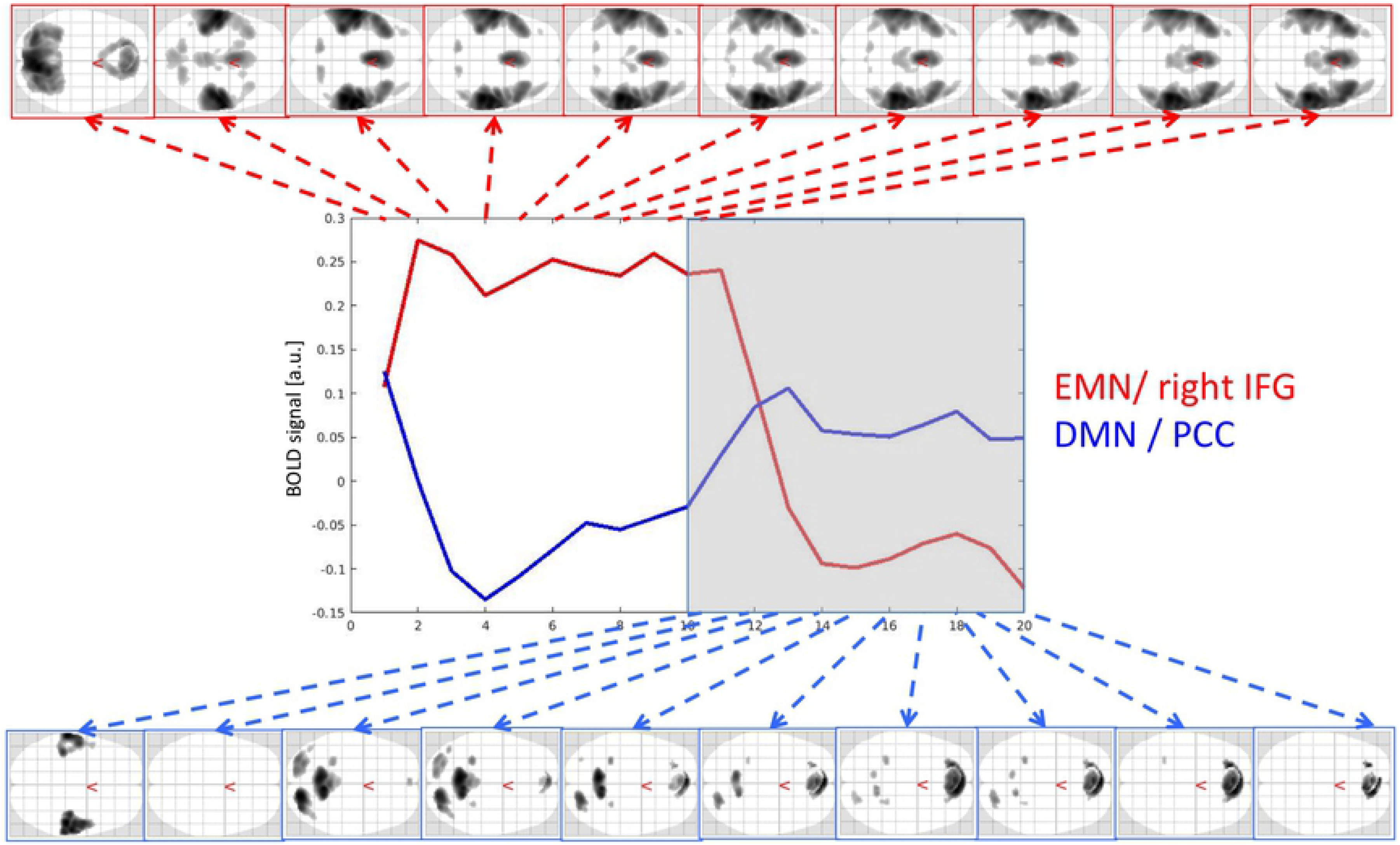
Figure 3 shows the corresponding axial glass brains for the ON- (upper row) and OFF-periods (lower row), and split for corresponding time-bins (TBs). The line-graphs in the middle of the figure show the development of the BOLD response across time for the ON (left side) and OFF (right side) period, respectively, and extracted from the posterior cingulate cortex (PCC) and inferior frontal gyrus (IFG), representing the default mode network (blue line, DMN) and extrinsic mode network (red line, EMN), respectively.

When the time-derivative (TD) contrasts were explored, which reflect a significant increase or decrease of activity from one time-bin to the next, activity changes were observed only at the transitions between the blocks. When the ON block started (see Figure 4), the EMN switched on at once, together with the visual response to the instructions on the screen.

**Figure 4:**
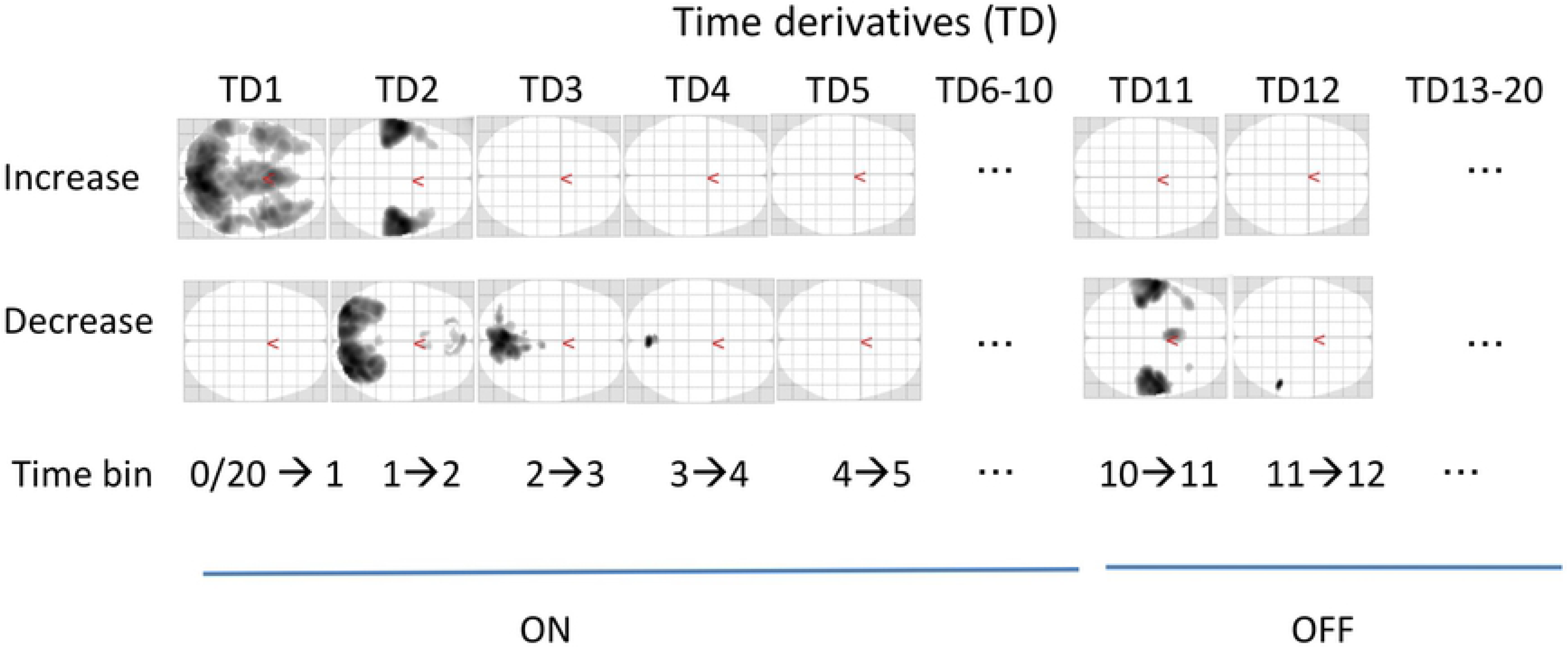
Figure 4 shows the axial glass brains obtained from the time-derivative (TD) analysis during ON (left side) and OFF (right side) periods, separately for each time-derivative contrast, contrasting time-bin (TB)1 with TB2, TB2 with TB3 etc in a “sliding window” through all TBs (1-20). The upper row of glass brains show increases in activity, the lower row shows decreases in activity. See text for further details.

However, the visual activity faded away over the first three TBs (cf. Logothetis et al., 2001), i.e. over the first 11-16.5 sec, while the auditory activity faded in from TB1 to TB2 and remained stable after that. At the end of the ON block, the auditory and articulatory motor activity rapidly disappeared in the transition from TB 10 to TB 11, with a remaining decrease into TB 12 within the right planum temporale. Interestingly, the DMN was not observed in these time-derivatives-like contrasts, indicating that the DMN did not show a sharp onset of activity at the beginning of the OFF block, comparable to the EMN did at the beginning of the ON block. Neither EMN, nor DMN showed a sharp decrease of activity in this analysis.

## Discussion

To sum up the main findings, the ON- and OFF-blocks, corresponding to active task-processing versus passive resting, produced two orthogonal, non-overlapping activity patterns (see Figure 2). As is obvious from Figure 2, while task-present epochs resulted in a more anterior activity pattern, with the SMA/preSMA and the auditory cortices as the dominant activity regions, the task-absent epochs resulted in a more posterior activity pattern, including activity in the precuneus, and the parietal lobules as the dominant regions. These activity patterns would partially correspond to the EMN and DMM networks, respectively [1] [20]. 2001). The FIR analysis of the network dynamics based on time-derivative contrasts in addition revealed that the EMN showed a relatively sharp onset of the up-regulation at the beginning of the ON-blocks, while there was a more gradual up-regulation of the DMN during OFF-blocks. As expected, the DMN was more active during the OFF-blocks (see Figure 3), but no abrupt change was seen in the time-derivative analysis at the ON-OFF block intersections (see Figure 4). In their meta-analysis of nine PET blood-flow studies [19]) found significant decreases for the active minus passive task condition in cortical areas that today would qualify as the “classic” areas for the default mode network. One interpretation of the decreases in the Schulman et al. [19] study is that task-processing may inhibit activity in areas that otherwise would be tonically activated in the absence of a task, as was suggested by Popa et al. [50]. These authors used electrophysiology recordings and found that local field potential power was lower in anterior cingulum and retrosplenial cortex during task-OFF compared to task-ON periods, while the reverse was found in somato-sensory association cortex and middle temporal gyrus. A further confirmation of this hypothesis would be if activity in approximately the same areas would be increased during passive resting periods in between active task processing periods, which the current results have shown. The areas showing decreased activity during active minus passive epochs in the Schulman et al. [19] study were primarily in the posterior cingulate/precuneus, dorsolateral and inferior frontal cortex, and in the inferior temporal gyrus (passive here meaning being exposed to the same stimulus, but without instruction to act on the stimulus). Approximately the same areas were activated in the present study, but now when subtracting active task-processing periods from activity during a resting period, which would confirm the hypothesis of inhibition of tonically active areas during phasic task-processing. Similarly, Schulman et al. [19] found increases in the visual cortex in the occipital lobe during active visual task processing, after averaging data from 10 different studies with visual tasks. This is paralleled in the present study which found corresponding increases in the auditory cortex in the temporal lobe to an auditory task. Previous studies have shown that the DMN is still up-regulated during task-presence periods but attenuated or suspended compared to task-related activity, e.g [51], [52], [10]. Although the present findings are in line with these results, we cannot say if the corresponding networks were down-regulated or merely attenuated during reversals. An advantage with the present paradigm over previous paradigms that have been used, e.g. [53], [2], [54] is that the three different cognitive tasks were all embedded within the same experimental paradigm. The dichotic listening (DL) task is moreover exceptionally easy to understand and perform, so that the understanding of the task in itself does not require the allocation of additional cognitive resources which could confound task processing.

Figure 3 shows that the time-course for the task-positive EMN network followed a square-like trajectory, with a rapid initial up-regulation during ON-periods which peaked latest after about 11 sec (TB2), followed by a similarly rapid down-regulation during OFF-periods, peaking after about 22 sec (TB14). The corresponding trajectory for the task-negative DMN network showed a similar rapid down-regulation during ON-periods, peaking after about 22 sec (TB4), while the up-regulation during OFF-periods were more gradual than for the EMN network, actually beginning already in the middle of the ON-period (see Figure 3). This could be an anticipatory effect of waiting for the “next task to be presented”, and may represent an anticipatory shift of attention focus from task processing to resting, a kind of “readiness” for what is to come [55], which previously has been associated with alterations in EEG alpha-activity [56], [57]. A look at the time-course trajectories in Figure 3 shows a marked difference in the “gap” between the up-regulated and down-regulated network for ON versus OFF blocks. Although the two networks to a certain extent were down-regulated at about the same level, the level of up-regulation for the EMN was about twice the level of up-regulation for the DMN, which caused a gap difference between the two networks. The gap difference could reflect the additional increase in metabolism demands during active task-processing compared to resting. There was finally a delay for EMN up- and down-regulation of about 5.5 sec, which may reflect the delay and slowness of the BOLD-response in itself [58], and not an effect of network interaction and interference.

**In conclusion**, the EMN and DMN networks seem to alternate with the same frequency as the switch from active task-periods (ON-blocks) to passive rest-periods (OFF-blocks) which alternated on a 55 sec basis. From recent work that the DMN may represent a unique state of the mind e.g. [59], [60], [61], [52], we now suggest that the DMN is a marker of an egocentric state of mind, while a task-positive network, like the extrinsic mode network (EMN) [1] is a marker of an allostatic state of mind. These states dynamically fluctuate over time, such that the individual is either in one or the other state, with corresponding network activity being dominant in a particular state, and that this is mapped onto how environmental processing demands change over time. It may finally be suggested that the dynamic interaction between task-positive and task-negative networks may be disrupted in certain psychiatric and neurological disorders, extending what previously has been suggested for DMN abnormality [62], [63], [64]. We now suggest that what is abnormal in certain mental disorders may not be so much abnormality of a single network, but rather abnormality of network interaction and network dynamics, and that this is better captured in an experimental design with alternating task-ON and task-OFF periods, rather than a prolonged resting period during the scanning e.g. [65], [66], [67].

## Acknowledgements

The authors want to thank Roger Barndon, Eva Øksnes, Turid Randa for running the MR scanner, and Merethe Nygård, Heidi van Wageningen, Kerstin von Plessen, and Margaretha Dramsdal for collecting the fMRI data that were used in the present study. We also thank Marcus E. Raichle, Washington University School of Medicine, USA, for constructive comments on earlier versions of the paper.

## Conflict of interest

The authors declare no conflict of interest. The co-authors KH, LE, RG and KS own shares in the company NordicNeuroLab Inc., which produced the headphones used for the presentation of the auditory stimuli. These authors declare no conflict of interest.

**The present study was supported by grants** to Kenneth Hugdahl from European Research Council (ERC AdG #693124), the Western Norway Health Authorities (Helse-Vest, #912045), and the Research Council of Norway (NORMENT #223273)

